# Long-term spatially-replicated data show no physical cost to a benefactor species in a facilitative plant-plant interaction

**DOI:** 10.1101/2022.10.17.512641

**Authors:** Morgan J. Raath-Krüger, Christian Schöb, Melodie A. McGeoch, Divan A. Burger, Tanya Strydom, Peter C. le Roux

## Abstract

Facilitation is an interaction where one species (the benefactor) positively impacts another (the beneficiary). However, the reciprocal effects of beneficiaries on their benefactors are typically only documented using short-term datasets. We use *Azorella selago*, a cushion plant species and benefactor, and a co-occurring grass species, *Agrostis magellanica*, on sub-Antarctic Marion Island, comparing cushion plants and the grasses growing on them over a 13-year period using a correlative approach. We additionally compare the feedback effect of *A. magellanica* on *A. selago* identified using our long-term dataset with data collected from a single time period. We hypothesized that *A. selago* size and vitality would be negatively affected by *A. magellanica* cover and that the effect of *A. magellanica* on *A. selago* would become more negative with increasing beneficiary cover and abiotic-severity, due to, e.g., more intense competition for resources. We additionally hypothesized that *A. magellanica* cover would increase more on cushion plants with greater dead stem cover, since dead stems do not inhibit grass colonization or growth. The relationship between *A. magellanica* cover and *A. selago* size and vitality was not significant in the long-term dataset, and the feedback effect of *A. magellanica* on *A. selago* did not vary significantly with altitude or aspect; however, data from a single time period did not consistently identify this same lack of correlation. Moreover, *A. selago* dead stem cover was not significantly related to an increase in *A. magellanica* cover over the long term; however, we observed contrasting results from short-term datasets. Long-term datasets may, therefore, be more robust (and practical) for assessing beneficiary feedback effects than conventional approaches, particularly when benefactors are slow-growing. For the first time using a long-term dataset, we show a lack of physical cost to a benefactor species in a facilitative interaction, in contrast to the majority of short-term studies.

## Introduction

Biotic interactions strongly shape ecological communities, with these interactions affecting plant fitness, abundance, and survival (Maestre et al. 2010, Montgomery et al. 2010, Llambí et al. 2018, Raath-Krüger et al. 2021). The impact of biotic interactions, however, can vary considerably both spatially (Maestre et al. 2010, Mod et al. 2016) and temporally (Holzapfel and Mahall 1999, Armas and Pugnaire 2009, le Roux et al. 2013), and can scale up to influence species’ richness, distributions, and community composition (Adler et al. 2012, Llambí et al. 2018, Raath-Krüger et al. 2019). Indeed, variation in the outcomes of biotic interactions is strongly linked to environmental conditions in many systems (Kawai and Tokeshi 2006, Zhang and Tielbörger 2020), with a higher prevalence of net positive interactions (e.g., facilitation) typically observed under greater environmental severity (as posited by the Stress Gradient Hypothesis SGH; Bertness and Callaway 1994). However, under moderate environmental conditions, negative interactions (e.g., competition) are predicted to dominate. Therefore, in areas of high environmental severity or at the extreme ends of environmental stress gradients, positive effects of facilitation by neighbours may outweigh the negative effects of competition (He et al. 2013; although see the concept of facilitation collapse: Michalet et al. 2014).

Plant-plant facilitation is a non-trophic interaction where at least one of the interacting species benefits (e.g., through increased survival and/or performance; Schöb et al. 2014a,b). However, while most research only documents the positive impact of one species (i.e., the benefactor species) on associated beneficiaries (Armas et al. 2011, Hupp et al. 2017, Zhang and Tielbörger 2020), the impacts of beneficiaries on their benefactors (i.e., beneficiary feedback effects) are sometimes ignored (although see Pugnaire et al. 1996, Michalet et al. 2011, Schöb et al. 2014a,b,c, García et al. 2016, Michalet et al. 2016, Llambí et al. 2018, Melfo et al. 2020). Studies that have documented the bidirectional (i.e., reciprocal) nature of interactions between benefactor and beneficiary species have found that beneficiary species can have positive (Pugnaire et al. 1996), neutral (Lortie and Turkington 2008) or negative (Cranston et al. 2012, Schöb et al. 2014b,c, García et al. 2016, Michalet et al. 2016, Llambí et al. 2018) effects on their benefactors. For example, Schöb et al. (2014c) documented lower seed set and flower density for the cushion plant *Arenaria tetraquetra* under greater beneficiary species cover. In contrast, Pugnaire et al. (1996) showed a mutualistic interaction between a benefactor shrub (*Retama sphaerocarpa*) and a beneficiary species, where both species displayed improved performance when co-occurring. Llambí et al. (2018) examined reciprocal effects between the cushion plant *Arenaria musciformis* and two other species: cushion plant flower density decreased with increasing native plant density, but the non-native plant species had an indirect positive effect on the cushion plants due to its negative impact on the density of the native species (see also Melfo et al. 2020). Reciprocal interactions within a facilitative system may therefore have both strong and diverse impacts, and thus an understanding of the bidirectional nature of biotic interactions is essential for accurately assessing the consequences of plant-plant interactions (Schöb et al. 2014a,b,c, Melfo et al. 2020).

To date, studies examining reciprocal plant-plant interactions that include facilitation have typically used data from short-term experiments (e.g., Holzapfel and Mahall 1999, Lortie and Turkington 2008, but see Metz and Tielbörger 2016) and/or observations from a single time period (i.e., snapshot data; e.g., Pugnaire et al. 1996, Llambí et al. 2018). Therefore, studies describing the long-term impact of biotic interactions in systems where facilitation is prominent are lacking (but see Armas and Pugnaire 2005, Miriti 2006, García et al. 2016). The approach of monitoring the same species and/or individuals using long-term datasets (i.e., through repeated measurements of the same interacting individuals, e.g., repeat photography) is an effective method for examining changes in community structure and composition (Magurran et al. 2010, Fitzgerald et al. 2021). However, there are limitations associated with these types of correlative methods, including their inability to demonstrate causality and to provide a mechanistic understanding of the link between abiotic conditions and changes in the outcome of plant-plant interactions (Metz and Tielbörger 2016). Monitoring approaches can, however, provide valuable insights into the long-term consequences of biotic interactions (e.g., Báez and Collins 2008, Ribeiro et al. 2018). For example, extended monitoring of interacting individuals can disentangle short- and long-term responses to a disturbance (e.g., experimental removal of individuals to examine the influence of competition; see Maestre et al. 2003, Kikvidze et al. 2006). Moreover, monitoring approaches may be especially valuable for examining interactions between slow-growing and/or long-lived perennial species (e.g., many species in abiotically-stressful environments; Armas and Pugnaire 2005) because these species may respond slowly to changes in their interactions with other species (e.g., Raath-Krüger et al. 2021). Monitoring approaches could therefore be used to accurately examine the reciprocity of biotic interactions.

The aim of this study is to examine the reciprocal feedback effect of a beneficiary species on its benefactor species, creating a more complete understanding of the impact and bidirectional nature of the interaction between these two plant species. We used a widespread cushion plant species and benefactor, *Azorella selago* (*Azorella* hereafter), and a dominant perennial grass species, *Agrostis magellanica* (*Agrostis* hereafter), on the sub-Antarctic Marion Island as a model system. We document the long-term outcome of the *Azorella-Agrostis* interaction by assessing changes in *Azorella* size and vitality in relation to *Agrostis* cover over a 13-year time period, and test if changes in the cover of *Agrostis* over the same period are related to changes in the characteristics of *Azorella* (i.e., size and stem mortality). We additionally compare these results with those obtained when using the more conventional snapshot approach (i.e., using data collected separately from a single time period – in 2003, 2006, and 2016; following, e.g., Pugnaire et al. 1996, Llambí et al. 2018). We hypothesize *Azorella* size and vitality would be negatively affected by *Agrostis* cover (due to, e.g., shading; see le Roux et al. 2005, and/or competition for resources; see Owen 1995), and that *Agrostis* cover would be positively related to *Azorella* dead stem cover (since dead stems do not inhibit grass colonization or growth while still providing the same benefits as live *Azorella* stems; see Michalet et al. 2011, Whinam et al. 2014).

We also examine the *Azorella-Agrostis* interaction across a gradient of environmental severity because the outcome of biotic interactions may shift in response to changing abiotic conditions (e.g., in line with the SGH: Kawai and Tokeshi 2006). Variation in the impact of *Azorella* on *Agrostis* has been documented along environmental gradients, with, for example, the impact of *Azorella* on *Agrostis* switching from negative to positive with increasing environmental severity across both fine and broad spatial extents (see le Roux and McGeoch 2010, le Roux et al. 2013). Additionally, plant populations on Marion Island may experience different abiotic conditions across the eastern and western sectors of the island (see, e.g., Goddard et al. 2021). However, no data are available to test if the reciprocal impact of *Agrostis* on *Azorella* varies with increasing abiotic stress. We hypothesise that beneficiary feedback effects will become more negative with increasing beneficiary cover and abiotic-severity (i.e., altitude), with the negative effect of *Agrostis* cover on *Azorella* growth and vitality being stronger under greater abiotic stress due to, e.g., more intense competition between *Azorella* and *Agrostis* for resources (following observations by Michalet et al. 2016, albeit in a different system).

## Material and Methods

### Study site and species

Marion Island is located in the southern Indian Ocean (46°55’S, 37°45’E), and has a hyper-oceanic climate (Smith and Steenkamp 1990), with low but very stable temperatures (mean annual temperature = 5.1 °C, mean diurnal range = 1.9 °C), along with high precipitation (2500 mm per annum) and humidity, cloud cover on most days and frequent strong winds (le Roux 2008). There is a clear elevational gradient of increasing abiotic severity on the island, with greater wind speeds and lower mean temperatures and soil stability at higher elevations (Boelhouwers et al. 2003, le Roux 2008). Consequently, plant species richness (le Roux and McGeoch 2008a, Chown et al. 2013), cover (Smith et al. 2001) and productivity (Smith 2008) decline with increasing elevation on the island. Moreover, the western and eastern aspects of Marion Island differ in terms of at least some abiotically conditions (see e.g., Goddard et al. 2021), resulting in plant populations across the two aspects potentially experiencing different abiotic stressors.

Marion Island supports 22 indigenous and 17 alien vascular plant species and over 200 bryophyte and lichen species (Greve et al. 2019, Kalwij et al. 2019). *Azorella selago* Hook. (Apiaceae) is a widespread cushion plant species (i.e., a compact, hemispherical species) occurring on multiple sub-Antarctic islands and in many habitat types on those islands. Due to its cushion growth form, *Azorella* ameliorates stressful environmental conditions (Nyakatya and McGeoch 2008, McGeoch et al. 2008), particularly in the cold, wind-exposed areas where the cushion plant is common. Consequently, *Azorella* hosts an array of species, including invertebrates and other plant species (Huntley 1971, Huntley 1972, Barendse and Chown 2001, Hugo et al. 2004), which makes the plant an important ecosystem engineer and keystone species (Hugo et al. 2004).

The demography of *Azorella*, and the biotic and abiotic factors driving performance and survival in the species, has not been extensively studied (but see le Roux et al. 2005). *Azorella* plant size and growth rate differ significantly between sites on Marion Island (Nyakatya 2006; McGeoch et al. 2008), and *Azorella* size is not related to altitude (McGeoch et al. 2008). In terms of vitality, *Azorella* experiences greater stem death and early senescence under reduced rainfall conditions, implying that the species may be negatively impacted by climate change (see le Roux et al. 2005, see also Frenot et al. 1997). However, cushion plant vitality may also be associated with a variety of other factors including climatic extremes (e.g., wind-exposure), competition for resources (i.e., due to increases in the cover and/or density of other vascular and non-vascular species on the cushion plants; see, e.g., Owen 1995), and mechanical damage from burrowing species and/or pathogens (Phiri et al. 2009; see also, Whinam et al. 2014).

*Agrostis magellanica* (Lam.) is a dominant perennial grass species on Marion Island and is the most common vascular plant species growing on *Azorella* (Huntley 1972). *Agrostis* occurs in most habitats on Marion Island and has the second largest altitudinal range (*c*. 0 - 600 m a.s.l.) of all the vascular plant species after *Azorella* (Huntley 1971, le Roux and McGeoch 2008a). *Agrostis* grows tall in wet manured areas but remains short in wind-exposed areas and is chiefly confined to the leeward side of *Azorella selago* cushions in fellfield habitats (van der Merwe et al. 2021). *Azorella* and *Agrostis*, together, form an appropriate model system for this study for several reasons. First, although *Azorella* can host dense and diverse epiphyte (Huntley 1972) and invertebrate communities (Hugo et al. 2004), *Agrostis* is by far the dominant epiphyte on *Azorella* and, in certain instances, covers > 50% of the surface of *Azorella*. Second, while the positive impact of *Azorella* on the local microenvironmental conditions and on *Agrostis* population structure and performance has been well-studied (see le Roux and McGeoch 2010, le Roux et al. 2013, Raath-Krüger et al. 2021), the reciprocal impact of *Agrostis* on *Azorella* has not yet been tested (although see Schöb et al. 2014a,b,c). Third, *Azorella* and *Agrostis* are characteristic species of the dominant vegetation type on Marion Island (and across the sub-Antarctic region) – the fellfield vegetation complex (Gremmen and Smith 2008) in which this study was conducted.

### Data collection

The outcome of the interaction between *Azorella* and *Agrostis* was inferred from changes in *Azorella* size and dead stem cover and *Agrostis* cover through time. Twelve long-term monitoring plots, which were established in 2003 at three altitudes (*c*. 200, 400 and 600 m a.s.l) on the island’s eastern and western aspects (Nyakatya and McGeoch 2008; see Supporting information for details), were resurveyed in 2006 and 2016 (plants were surveyed and photographed in these three years only due to the isolated and relatively inaccessible nature of the study site). One plot was positioned within each altitudinal band, forming two altitudinal transects (with each altitudinal transect comprising three plots) on the eastern (NE and SE transects) and western (NW and SW transects) sectors of the island (Supporting information). Fifty *Azorella* individuals were surveyed from each of the plots in each altitudinal band, with a total of n = 300 *Azorella* individuals surveyed from both the eastern and western aspects of Marion Island (i.e., 600 *Azorella* individuals were surveyed in total across the island). Each *Azorella* individual was photographed in the summer of 2002/2003 (Nyakatya 2006, Nyakatya and McGeoch 2008) from directly above at a height of 1.5 m, with a scale bar included within each photograph. Each *Azorella* individual was photographed again in the summers of 2006 and 2016 using the same methods (see Supporting information).

To assess how *Azorella* size and stem mortality changed in relation to *Agrostis* cover, the photographs for each year were analysed in Fiji ImageJ (Schindelin et al. 2012) and Adobe Photoshop (Adobe Photoshop CS 2004). *Azorella* size and dead stem area were measured using the polygon area selection tool, the wand tracing tool and by adjusting colour thresholds in Image J (Supporting information). *Azorella* size was defined as the total horizontal surface area of the cushion plants (including live stems, dead stems enclosed by live stems, dead stems contiguous with live stems, and parts of the cushion plants that were covered by other plants) as observed from directly above in the photographs. The extent of dead stems was used as a measure of plant vitality, where higher cover of dead stems on a plant was assumed to represent lower vitality (following Huntley 1972). The area of *Agrostis*, other vascular plants, non-vascular plants and rock were measured using the same methods (Supporting information). Where other vascular and non-vascular plant species were growing on the edge of cushion plants and the edges of the cushion plants were not directly visible, we interpolated the plant edge based on the shape of the cushion plants. Similarly, we interpolated the cover of other vascular plant species and mosses when their cover was obscured by *Agrostis* cover; however, this was rare. In almost all cases, the vascular and non-vascular plants measured on the *Azorella* individuals were alive. Generally, very few dead individuals of other species are observed growing on *Azorella*, likely due to strong wind on Marion Island rapidly removing dead individuals. Dead stems were defined as any portions of *Azorella* individuals that were black or grey (and/or had very low stem densities) and/or if *Azorella* stem tips consisted entirely of brown leaves (following Bergstrom et al. 2015; this did not include autumnal senescence due to photographs having been taken in mid- or late-summer).

Due to windiness during photography and the unevenness of the substrate there was some random variation in the quality of the photographs: of the 1500 images that were analysed, 414 images were excluded from the dataset because: 1) the images were of poor quality (e.g. snow or mist in the photographs limited the accuracy of the measurements); 2) the images were taken at slightly different angles; 3) the incorrect individuals were re-photographed in 2016; or 4) *Azorella* individuals died or only small fragments of the individuals remained. One additional image was removed from the dataset as an outlier because the *Azorella* individual had a very high cover of non-vascular species (i.e., 31% cover, where on average non-vascular plant species cover on *A. selago* was 1.0 ± 0.1% [mean ± SE] in 2003, 1.8 ± 0.2 in 2006, and 2.1 ± 0.1% in 2016). Only cushion plant individuals with suitable photographs from both 2003, and 2016 were included in the final analyses (i.e., of the 600 cushion plants surveyed, 445 were included in the analysis). While images from all of the sites were processed for 2003 and 2016, for 2006 a subset of images (i.e., from one easterly and one westerly transect) were processed (of the 300 images analysed in 2006, 196 images were suitable for inclusion in the analysis).

### Data analysis

To test if *Agrostis* had a long-term negative effect on *Azorella* growth, a generalized linear mixed model (GLMM) was used to model final *Azorella* size (i.e., *Azorella* area in 2016) in response to initial *Azorella* size (i.e., cm^2^ in 2003), initial *Agrostis* cover on *Azorella* (i.e., % cover), initial *Azorella* dead stem cover (%), initial cover of other vascular plants and mosses on *Azorella*, aspect, and altitude (Supporting information). To examine whether *Agrostis* negatively impacts *Azorella* vitality, a second GLMM was run using final *Azorella* dead stem cover as a response variable. The time-series nature of the data enabled us to test whether *Agrostis* cover in the past affected later *Azorella* performance. Finally, because dead *Azorella* stems are less densely packed than live *Azorella* stems, and dense cushion plants can have a lower epiphyte cover than lax cushions (Bonanomi et al. 2016, Jiang et al. 2018) and because *Agrostis* has been shown to have a higher cover on the dead parts of a congeneric cushion plant species on Macquarie Island (i.e., *Azorella macquariensis*; see Whinam et al. 2014), we assessed whether dead *Azorella* stems facilitated an increase in *Agrostis* cover through modelling final cover of *Agrostis* on *Azorella* in response to initial *Agrostis* cover, initial *Azorella* size, initial *Azorella* dead stem cover, initial cover of other vascular plants and mosses on *Azorella*, aspect, and altitude (Supporting information). The same models were repeated using the data surveyed in 2006 to examine the short-term impact (i.e., between 2003 and 2006) and medium-term impact (between 2006 and 2016) of the *Azorella-Agrostis* interaction.

The initial cover of other vascular plants and mosses was included in all models to account for the effect of these sub-ordinate species. To account for the data’s spatial structure, a random effect for “plot” was included in each model. All *Azorella* individuals from the high-altitude plots were excluded from model assessing the influence of initial *Azorella* dead stem cover on final *Agrostis* cover since *Agrostis* was absent from most of these individuals (*c*. 95%) in both the final and initial year of the survey. Two interaction terms were included in each model: 1) ‘initial *Agrostis* cover × altitude’ and ‘initial *Agrostis* cover × aspect’, since altitude and aspect may mediate the effect of *Agrostis* cover on *Azorella* size and vitality; and 2) ‘initial *Azorella* dead stem cover × altitude’ and ‘initial *Azorella* dead stem cover × aspect’, since the effect of *Azorella* dead stem cover on *Agrostis* cover may depend on altitude and aspect.

Final *Azorella* size (after log-transformation) was assumed to be normally distributed, whereas the percentage-type outcomes (i.e., final *Azorella* dead stem cover and final *Agrostis* cover were assumed to be beta distributed. The beta distribution is an appropriate distribution for the analysis of double-bounded data such as percentage data (Niekerk et al. 2019; see also, le Roux et al. 2013), and the logit link function was used to specify the relationship between the linear predictors and the mean percentages (Supporting information).

To reproduce the analyses that are typically used to examine beneficiary feedback effects, we additionally ran complementary GLMMs to examine the response of *Azorella* size, *Azorella* dead stem cover and *Agrostis* cover against predictor variables based on data collected from a single time period (i.e., 2003, 2006, and 2016; hence, using a snapshot approach; following e.g., Pugnaire et al. 1996, Llambí et al. 2018). For these analyses, we only included the same cushion plants that were used in the first analyses to allow a clear comparison between the two approaches.

Finally, analyses of *Azorella* growth were complemented with analyses based on the snapshot approaches of two additional measures of *Azorella* performance: the number of flower buds and the number of fruits. These data were available only for 2003 (from Nyakatya 2006). A GLMM based on the zero-inflated negative binomial distribution (to account for overdispersion and excess zeros) was used to determine whether *Agrostis* density was negatively related to (i) the number of flower buds and (ii) the number of fruits on *Azorella*. The model included “plot” as a random effect, and *Azorella* size was included in each model as an offset variable.

All analyses were conducted in R version 3.5.1 (R Core Team 2018) using the packages “car” (Fox and Weisberg 2011) and “glmmTMB” (Brooks et al. 2017).

## Results

Eighteen of the 600 *Azorella* individuals died over the 13-year observation period (i.e., annual mortality rate of *c*. 0.2% p.a. for plants > 15 cm diameter). The mean *Agrostis* cover on these 18 individuals (mean ± SE initial *Agrostis* cover: 2.6 ± 1.0 % in 2003) was less than that of the surviving *Azorella* individuals (mean ± SE initial *Agrostis* cover: 6.6 ± 0.5% in 2003). *Agrostis* cover increased on 58.3% of the 445 *Azorella* individuals (only considering plants with suitable photographs from 2003 and 2016; see Supporting information). Across all cushion plants, *Agrostis* cover on *Azorella* increased from 13.7 ± 1.1 % in 2003 to 18.0 ± 1.0 % in 2016 at the low altitude sites, and from 5.04 ± 0.6% to 6.85 ± 0.7% at the mid altitude sites (*Agrostis* was absent from the majority of *Azorella* individuals at the high-altitude sites in both years). The majority of *Azorella* plants (92%; Supporting information) increased in size, while *Azorella* live stem area increased on average from 1543.5 ± 59.2 cm^2^ in 2003 to 2044.5 ± 63.8 cm^2^ in 2016. Dead stem cover also increased on most (82%) of the *Azorella* plants (from 20.1 ± 0.6% in 2003 to 24.9 ± 0.6% in 2016).

In contrast to the very slow vertical growth rate of 0.43 ± 0.01 cm yr^−1^ (mean ± SE) of *Azorella* (le Roux and McGeoch 2004), cushion plant diameter increased by 1.98 ± 0.07 cm yr^−1^. Measurements of *Azorella* size and *Agrostis* cover from the images were highly correlated with the size measurements of the same individuals taken in the field, i.e., for circumference (*r*^2^ = 0.87 in 2003 and 0.84 in 2016), maximum diameter (*r*^2^ = 0.90 in 2003 and 0.89 in 2016), and *Agrostis* cover (*r*^2^ = 0.95 in 2016; *Agrostis* cover was not estimated in the field in 2003). Each of the reported correlations was statistically significant.

*Azorella* size was not significantly related to *Agrostis* cover in the multivariate models (Table 1). Additionally, initial *Agrostis* cover did not alter the relationship between the (i) final and initial *Azorella* size (Fig. 1a) and (ii) final and initial *Azorella* dead stem cover (Fig. 1b). *Azorella* dead stem cover, however, was significantly related to the interaction between initial *Agrostis* cover and aspect. Specifically, the positive relationship between initial *Agrostis* cover and final *Azorella* dead stem cover was stronger on the eastern side of Marion Island than on the western side. However, even though this interaction term was significant, its biological relevance was negligible as the difference between the slope estimates was small (i.e., slopes of 0.304 and 0.276, respectively, for the eastern and western aspects) and a comparison of AIC values showed that the model including the interaction term did not improve the model fit. More generally, *Azorella* size did not vary significantly with altitude or aspect; however, *Azorella* vitality was related to altitude, where dead stem cover was greater at low- and mid-altitudes compared to high altitudes (Table 1; Supporting information).

**Table 1.**
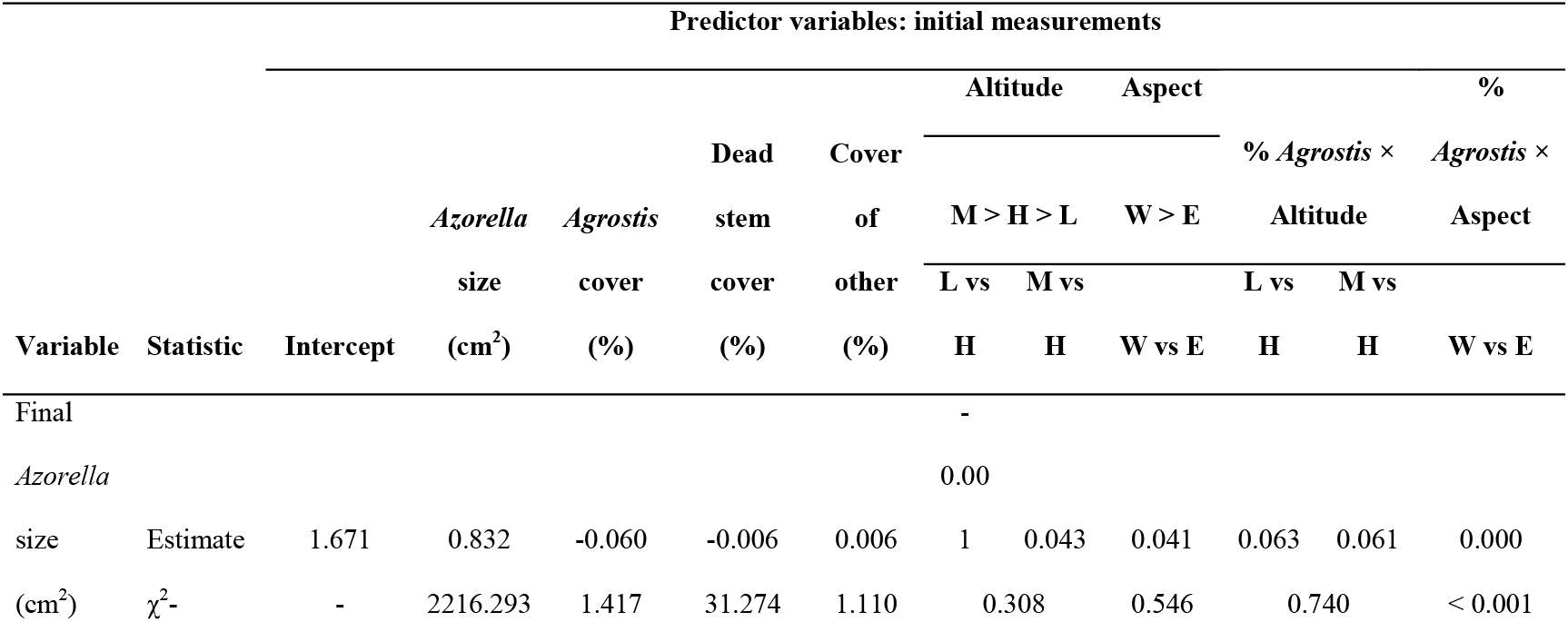
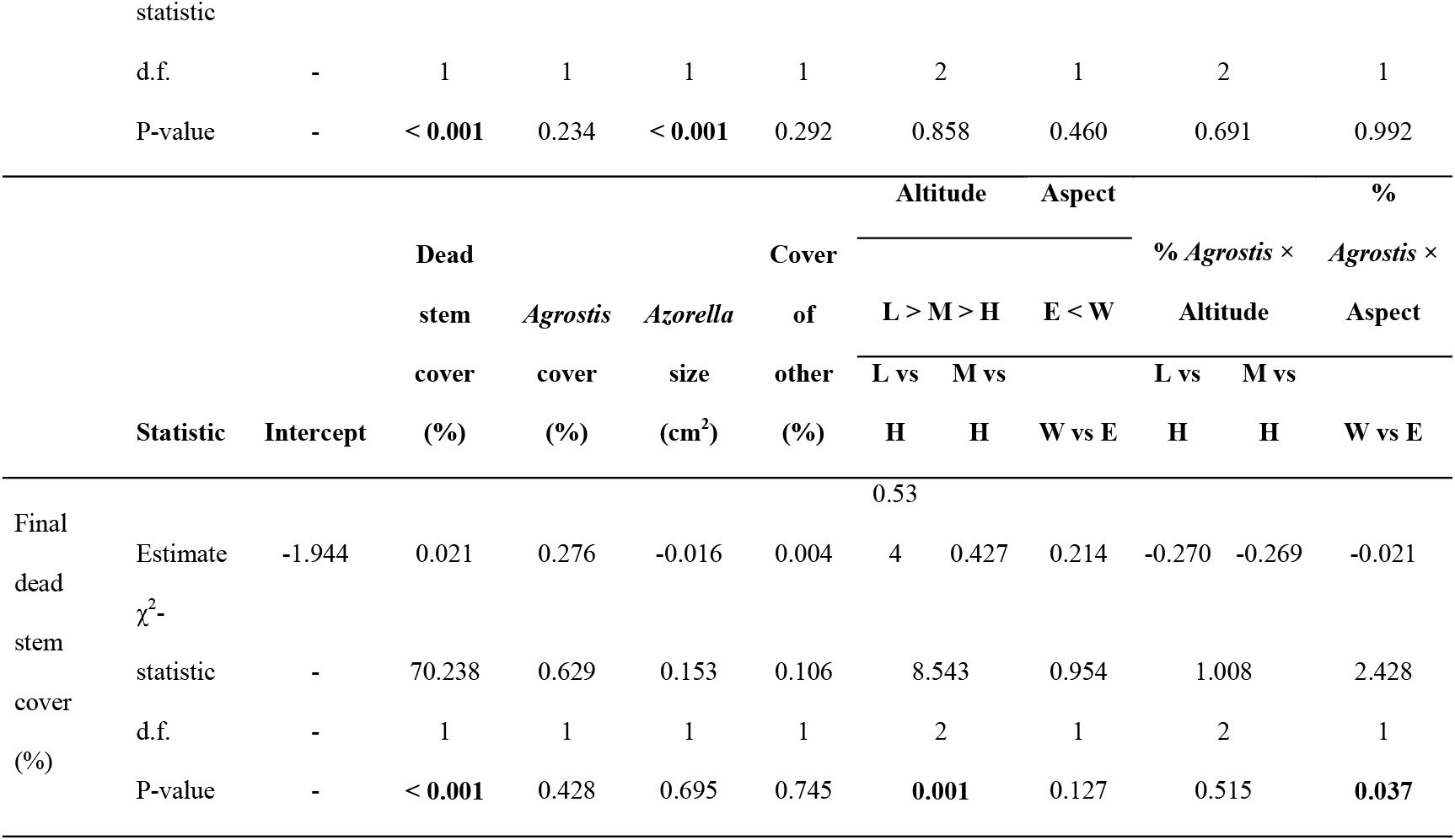
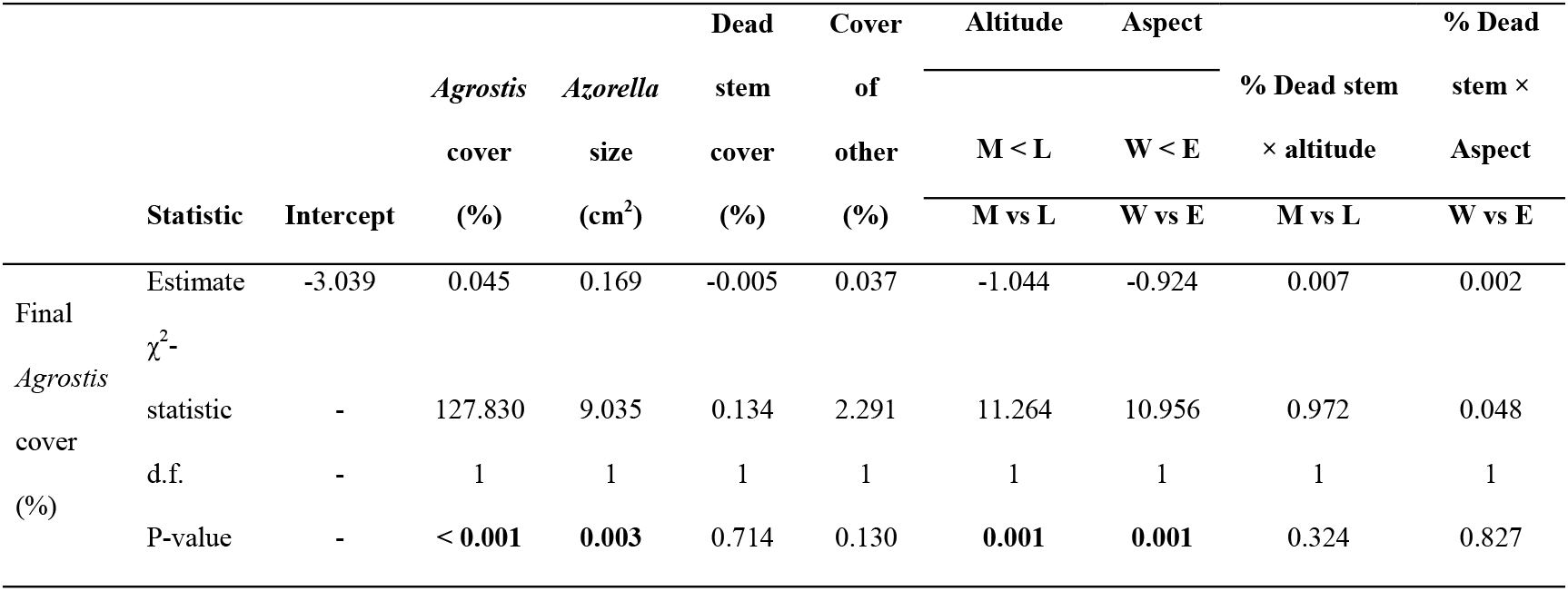
Modelling *Azorella selago* size (*n* = 445), *A. selago* dead stem cover (*n* = 445), and *Agrostis magellanica* cover (*n* = 317) between 2003 and 2016 using generalized linear mixed-effects models. Cover of other = combined cover of other vascular plant species and mosses. For both categorical variables (altitude and aspect), the factors levels’ are presented according to the order of their magnitude: L = Low, M = Mid, H = High, E = East, W = West. “L vs H”, “M vs H”, and “W vs E”, respectively, signify the difference between (i) low and high altitudes, (ii) mid and high altitudes, and (iii) western and eastern sides.

**Figure 1.**
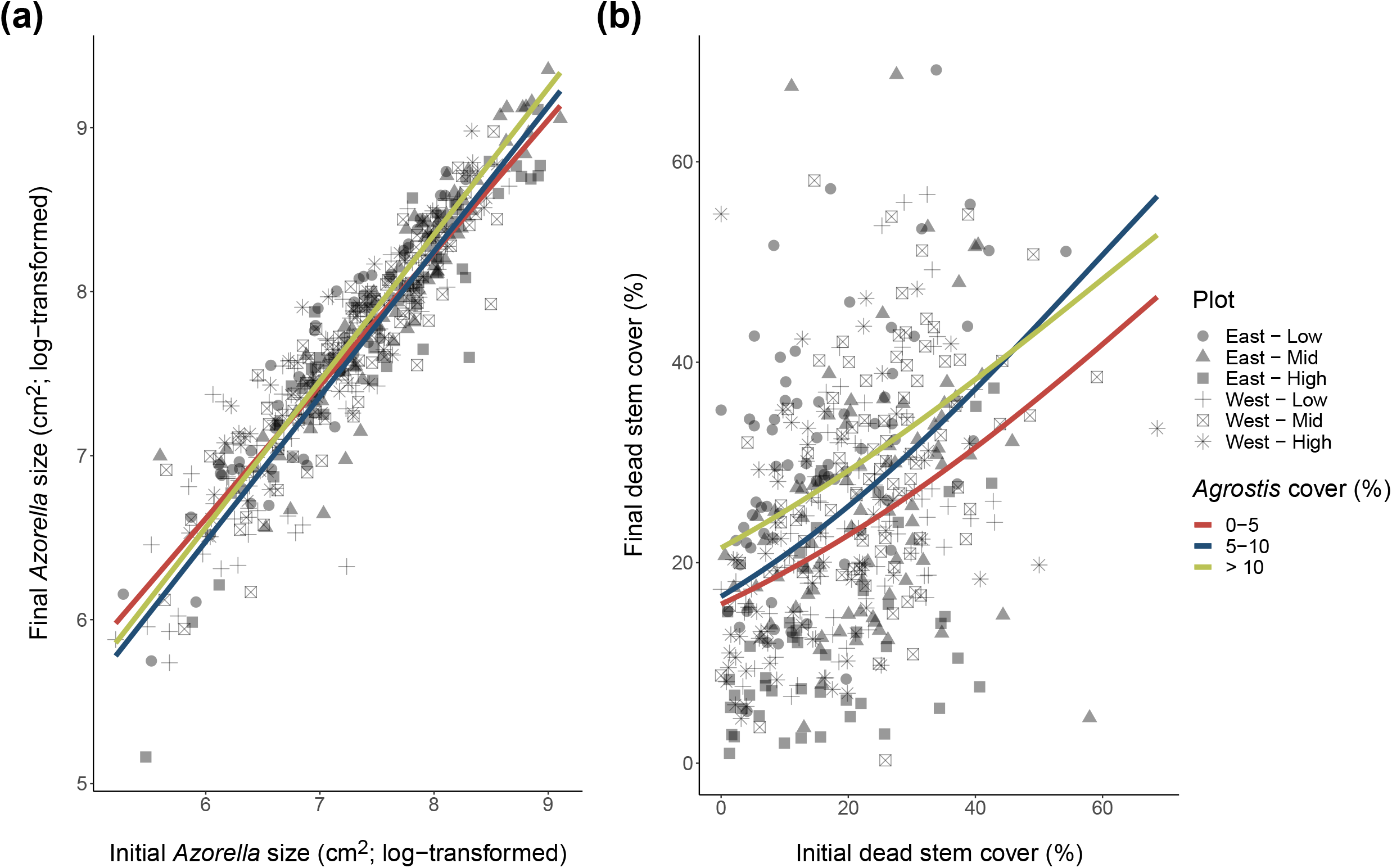
The relationship between a) final and initial *Azorella selago* size and b) final and initial *Azorella selago* dead stem cover with separate regression lines for three categories of initial *Agrostis magellanica* cover (i.e., 0-5%, 5-10 % or > 10 %; as measured in 2003). Symbols represent the sectors of the island (East or West) on which *A. selago* individuals were located and the altitude (low: c. 200 m a.s.l., mid: c. 400 m a.s.l. and high: c. 600 m a.s.l.) at which each individual occurred. The regression model showed that a) the relationship between final and initial *A. selago* size did not differ significantly between the initial *A. magellanica* cover categories (χ^2^-statistic = 5.236; d.f. = 2; P-value = 0.073), and b) the relationship between final and initial *A. selago* dead stem cover did not differ significantly between the *A. magellanica* cover categories (χ^2^-statistic = 0.485; d.f. = 2; P-value = 0.785).

Although the positive relationship between final and initial *Agrostis* cover was not affected by *Azorella* dead stem cover (Fig. 2), altitude and aspect were significantly related to final *Agrostis* cover (Table 1). Final *Agrostis* cover was significantly higher (i) at low altitudes compared to mid altitudes (low: mean ± SE = 17.97 ± 1.0 % vs mid: 6.85 ± 0.7 %) and (ii) on the eastern side of Marion Island compared to the western side (east: mean ± SE = 15.02 ± 0.9 % vs west: 3.75 ± 0.4 %; Table 1, Supporting information). Moreover, the relationship between final *Agrostis* cover and initial *Azorella* size was significantly positive, with larger *Azorella* individuals in the initial year having greater *Agrostis* cover in the final year. There was no significant effect of *Azorella* dead stem cover on *Agrostis* cover across 13 years (i.e., between 2003 and 2016). The results based on the analyses examining the short-term impact (i.e., between 2003 and 2006) and medium-term impact (between 2006 and 2016) of the *Azorella-Agrostis* interaction did not differ considerably, with only the sign and the significance of one term relating to our key hypotheses differing between the models (Supporting information). However, the analysis between 2006 and 2016 revealed that *Agrostis* cover decreased significantly with increasing *Azorella* dead stem cover (Supporting information).

**Figure 2.**
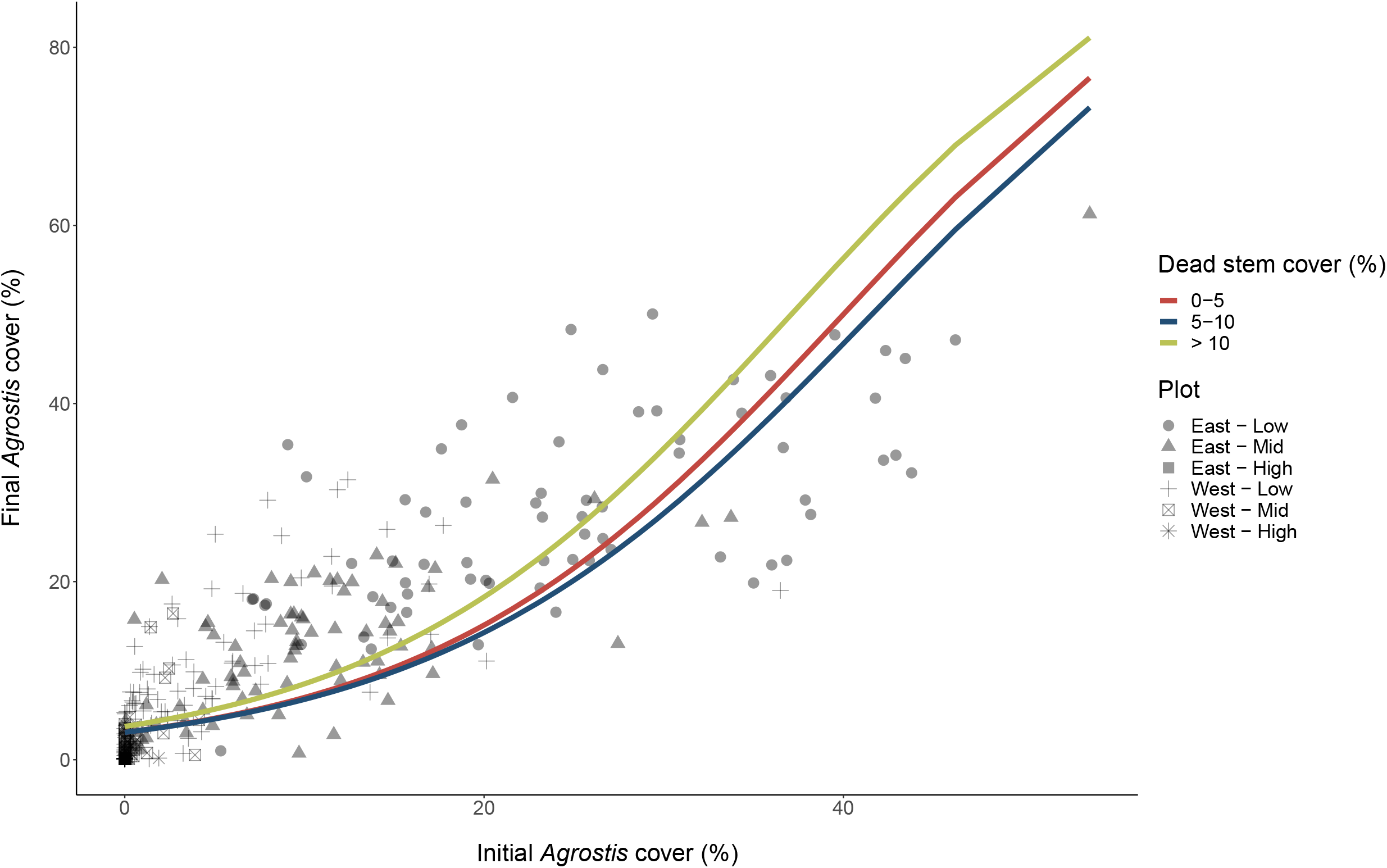
The relationship between final and initial *Agrostis magellanica* cover with individual regression lines drawn for three categories of *Azorella selago* dead stem cover (i.e., 0-5%, 5 10 % or > 10 %; as measured in 2003). Dead stem cover was calculated as: [(dead stem area in 2003/*Azorella* size in 2003) × 100]. The beta regression model showed that the relationship between final and initial *A. magellanica* cover did not differ significantly between the initial *A. selago* dead stem cover categories (χ^2^-statistic = 0.332; d.f. = 2; P-value = 0.847).

We observed contradictory results between our repeated measures approaches and the analyses based on data from a single time period, with the analyses based on data from a single time period showing different relationships between years for different measures of *Azorella* performance (Supporting information). Additionally, data from a single time period suggested that *Agrostis* cover was significantly positively related to the number of fruits on *Azorella* (Supporting information). In contrast, the number of flower buds on *Azorella* was significantly negatively related to *Agrostis* cover. The number of flower buds on *Azorella* was also significantly related to both aspect and altitude, with more flower buds on the western side of Marion Island than the east and fewer flower buds on *Azorella* at low and mid altitudes compared with high altitudes. In the snapshot analyses using data from 2003, 2006, and 2016, both *Azorella* size and *Azorella* dead stem cover were positively (and sometimes significantly) correlated with *Agrostis* cover (Supporting information).

## Discussion

In plant communities, the feedback effect of beneficiaries on their benefactors is infrequently considered and is typically only examined using short-term datasets and/or correlations over a single time period (see Pugnaire et al. 1996, Schöb et al. 2014a, Michalet et al. 2016). Here we used a long-term dataset of repeated measurements to examine the consequences of an interaction between a pair of plant species characterizing a dominant sub-Antarctic habitat. We made three main observations: 1) there was no significant effect of beneficiary species’ cover on benefactor size and vitality when using a long-term repeated measures approach, and the feedback effect of the beneficiary species did not vary with increasing abiotic stress, i.e., altitude, or with aspect (in contrast to, e.g., Michalet et al. 2016), 2) data from a single time period (i.e., using a snapshot approach) do not consistently reveal this same lack of correlation, and 3) the relationship between benefactor dead stem cover and beneficiary cover was not significant.

Our results revealed no significant long-term positive or negative relationship between *Agrostis* cover and *Azorella* size and vitality. Instead, the influence of *Agrostis* on *Azorella* may be negligible (in agreement with findings from some other benefactor-beneficiary systems; Armas and Pugnaire 2005, Lortie and Turkington 2008). Therefore, despite evidence for *Azorella* altering *Agrostis* population structure and increasing *Agrostis* biomass, reproductive output and abundance compared with surrounding areas where *Azorella* is absent (le Roux et al. 2013, Raath-Krüger et al. 2021), *Agrostis* has no significant reciprocal impact on cushion plant growth, vitality, and survival. Our results, therefore, suggest that the *Azorella-Agrostis* interaction could be considered commensalistic (see, e.g., Bronstein 2009).

We acknowledge that our correlative approach has methodological limitations, including its inability to reveal causal relationships (Metz and Tielbörger 2016). For example, microhabitat differences may cause covariation between *Azorella* and *Agrostis* growth (although no data exists to support this explanation). Additional experimental approaches may, therefore, be valuable because they provide a mechanistic understanding of the relationship between interacting species and the abiotic environment (Metz and Tielbörger 2016, Schöb et al. 2014b). Long-term monitoring approaches may still, however, be more beneficial than approaches using short-term and/or experimental datasets or data from a single time period (i.e., snapshot data) when examining interactions between slow-growing and/or long-lived perennial species (Armas and Pugnaire 2005) because these species may respond slowly to changes in their interactions with other species (Raath-Krüger et al. 2021). Specifically, on Marion Island, a repeated measures approach is more practical, and possibly more robust, for assessing beneficiary feedback effects than an experimental and/or snapshot approach because *Azorella* is so slow-growing (le Roux and McGeoch 2004) and may, therefore, respond gradually over longer periods to changes in biotic interactions and/or disturbances. Because cushion plants are long-lived organisms and can survive for several hundreds of years (Morris and Doak 1998), shorter-term differences in cushion plant size, dead stem cover and/or reproduction could be compensated for by changes in survival. However, given that it is impossible to study these plants over a lifetime, examining the performance of these plants from long-term observational datasets may provide deeper insights into the long-term balance of plant interactions, habitat dynamics and the evolutionary consequences for the species involved compared with datasets examining plant-plant interactions from a single time period (see, e.g., Schöb et al. 2014a,b, Barraclough 2015, Melfo et al. 2020).

Moreover, observational studies can be conducted at larger spatial scales relative to experimental and/or snapshot studies, providing increased potential for generalization and documentation of broad-scale patterns in plant communities (Stricker et al. 2015). By using an observational approach, we were able to examine approximately the full range of environmental severity (including the entire range of *Agrostis* cover on *Azorella*) within fellfield vegetation on Marion Island. Our results, therefore, highlight the value of using long-term datasets (in agreement with Metz and Tielbörger 2016) and suggest that to accurately assess the reciprocity of biotic interactions, observations from snapshot approaches examining the bidirectional nature of biotic interactions should be explicitly combined with long-term observational and/or long-term experimental approaches.

Contrary to expectation, *Agrostis* cover was not positively related to *Azorella* dead stem cover over the long term. However, the analysis based on the intermediate time period (i.e., between 2006 and 2016) revealed that final *Agrostis* cover decreased significantly with increasing *Azorella* dead stem cover, suggesting that *Azorella* cushion plants with a large cover of dead stems may not necessarily favour an increase in *Agrostis* cover. Therefore, changes in *Agrostis* cover may be linked to *Azorella* compactness (with greater *Azorella* dead stem cover leading to lower compactness), with compact cushion plants typically being associated with greater species cover, biomass, and richness, possibly due to higher substrate nutrient content, more effective heat-trapping and greater stability than lax cushion plants (Jiang et al. 2018, Yang et al. 2017, but see Al Hayek et al. 2015, Bonanomi et al. 2016). However, the opposite effect of compactness has also been observed in some cushion species (Michalet et al. 2011). For example, *Agrostis* grows easily on the dead parts of a congeneric cushion plant species, *Azorella macquariensis*, on Macquarie Island (Whinam et al. 2014). *Agrostis* cover did, however, increase significantly with *Azorella* size, suggesting that larger cushion plants may have a stronger positive effect on beneficiary cover compared to smaller cushion plants (in agreement with Yang et al. 2017). Moreover, final *Agrostis* cover varied significantly with altitude and aspect, with *Agrostis* cover being significantly higher on the eastern vs western side of Marion Island and at low altitudes compared to mid altitudes. Spatial variation in *Agrostis* cover on *Azorella* may be strongly linked to variation in the climate across Marion Island (le Roux and McGeoch 2008b, le Roux and McGeoch 2010). For example, western sectors of Marion Island are likely to be more abiotically-stressful than the eastern sectors because the western side of the island tends to experience stronger winds (Goddard et al. 2022). Moreover, plant species on Marion Island are exposed to increasingly stressful abiotic conditions (e.g., lower temperatures and greater wind speeds) with increasing altitude (see le Roux 2008). Consequently, the higher *Agrostis* cover observed on *Azorella* on the eastern side of the island may be due to spatial variation in abiotic stress across the island.

By using a snapshot approach, a broader range of benefactor performance metrics could be examined in this study. However, these results contrasted with some of the results from the repeated measures approach (i.e., from a long-term dataset, and from the short-term [i.e., 2003-2006] and medium-term [i.e., 2006-2016] datasets). In other words, measurements from a single time period can give different results depending on the timing of the measurements (in agreement with Trinder et al. 2013). For example, in this study, the snapshot approach suggests that the relationship between *Agrostis* and *Azorella* is neutral at one time, but positive or negative at another time. Some degree of the discrepancy between the long-term repeated measures approach and the snapshot approach could be explained by a shift in the outcome of the *Azorella-Agrostis* interaction over time (see e.g., Schöb et al. 2012). Indeed, abiotic conditions are important determinants of the variation in the outcome plant-plant interactions (Michalet et al. 2016). However, over the course of this study, there were minimal changes in temperature and rainfall (Shangheta 2020) and, due to the hyper-oceanic nature of the Marion Island’s climate, there is inherently minimal intra-annual variation in climate. It is, however, possible that long-term warming (i.e., over many decades) may contribute to an overall increase in beneficiary cover in the future, shifting the outcome of the benefactor-beneficiary interaction. For example, if climatic conditions become more benign (e.g., through warming temperatures), the growth rate of the beneficiary species may increase disproportionately quickly (relative to the benefactor species), and this may limit the performance of the benefactors (le Roux et al. 2005, Schöb et al. 2014b,c). Thus, while it has been speculated that the effect of benefactors on beneficiaries could shift under changing environmental conditions (see Armas et al. 2011), it is equally important to consider how the feedback effects of beneficiaries on their benefactors could change with shifting environmental conditions (e.g., Lin et al. 2012; Schöb et al. 2014a, García et al. 2016, Michalet et al. 2016). Relatedly, shifts in climatic conditions may indirectly result in a change in beneficiary cover as the outcome of interactions shift (e.g., as the intensity of the facilitative effect of the benefactor changes; Schöb et al. 2014a,b), and this can modify how the benefactor and beneficiary species interact (e.g., due to niche similarity effects, in which interactions of species sharing more similar niches with the benefactor are more competitive; Adler et al. 2012). However, a more complete understanding of the factors mediating the outcome of these interactions is required.

Second, the long-term outcome of plant-plant interactions likely varies with the beneficiary individuals’ ontogenetic stages (Eränen and Kozlov 2008, le Roux et al. 2013). For example, as *Agrostis* individuals grow, the effects of *Agrostis* on *Azorella* are likely to become more negative due to decreasing space or resources. However, these types of responses are not captured by the snapshot approach because snapshot approaches show the potential influence of beneficiaries on their benefactors at a specific point in time. In contrast, a repeated-measures approach is able to demonstrate the long-term effects of prevailing plant-plant interaction accumulated over time and provide insights into vegetation dynamics (Schöb et al. 2012; Fitzgerald et al. 2021). However, a greater understanding of the context-dependency of the long-term impacts of beneficiary feedback effects, as well as the generality of our findings, is needed (see also Schöb et al. 2014a).

In conclusion, while previous studies have reported negative impacts of beneficiaries on benefactor performance (Michalet et al. 2011, Schöb et al. 2014c), we show a lack of physical cost to a benefactor species in a facilitative interaction (i.e., at least over more than a decade). Moreover, the methodology used to examine feedback effects is important, as it may affect the observed outcome of biotic interactions (in agreement with Metz and Tielbörger 2016). A better understanding of the accuracy of the snapshot approach needs, therefore, to be determined and compared, or combined with, long-term repeated measures approaches and/or experimental approaches. The apparent contradiction between results also highlights the potential importance of long-term datasets in assessing the reciprocity of biotic interactions. Our findings suggest that studies examining beneficiary feedback effects need to move beyond just using contemporary snapshot approaches and/or short-term experiments because reciprocal interactions are dynamic, and considering temporal dynamism between interacting species may be key in understanding species co-existence (see e.g., Trinder et al. 2013, Schofield et al. 2018).

