## Supporting information for "Long-term spatially-replicated data show no physical cost to a benefactor species in a facilitative plant-plant interaction"

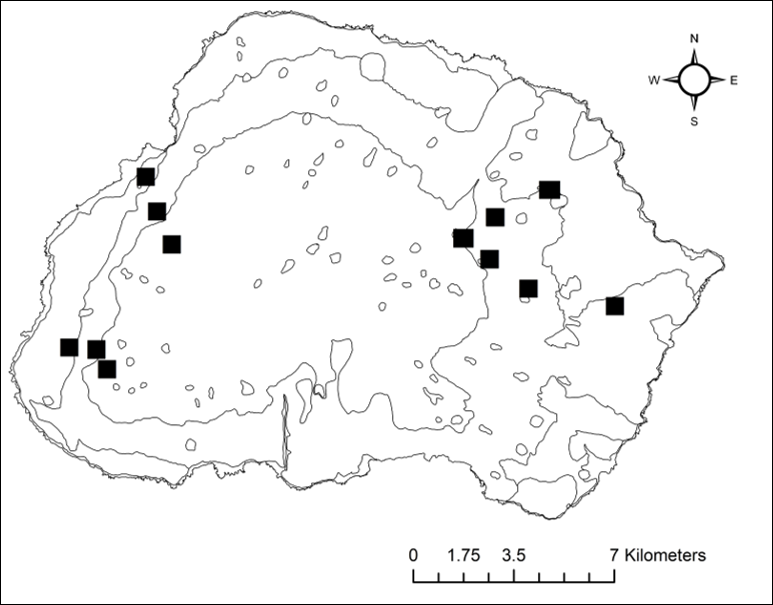

Supplementary Figure 1**.** Locations of the 12 plots (black squares) split into four altitudinal transects on sub-Antarctic Marion Island (see Supplementary Text S1 for details).

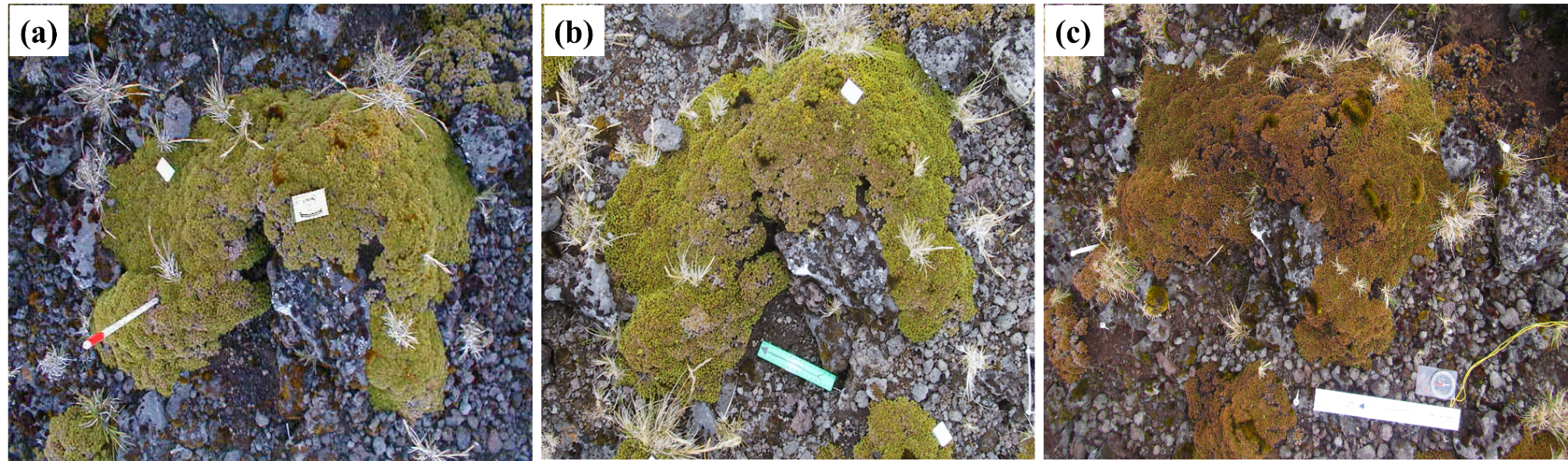

Supplementary Figure 2**.** Pictures of the same Azorella selago individual photographed in 2003 (a), 2006 (b), and 2016 (c) respectively, at the low altitude site on the western side of Marion Island. Photographs were taken directly from above with a scale bar (5.2 cm length matchbox, 15 cm and 30 cm length rulers) included. Digital cameras were used in 2003 (Nikon E885), and 2006 (Canon PowerShot S10) and in 2016 (Canon PowerShot D30). There was a difference in the resolution of the images taken between the years (300 dpi in 2003, and 180 dpi in 2006 and 2016); however, this did not affect the measurements as image processing was not performed at the highest resolution. Both cameras had standard lenses, which created minimal distortion, and because cushion plants were always photographed in the centre of the images, any distortions were negligible. Possible causes of A. selago damage on Marion Island include wind, alien house mouse burrowing and pathogens.

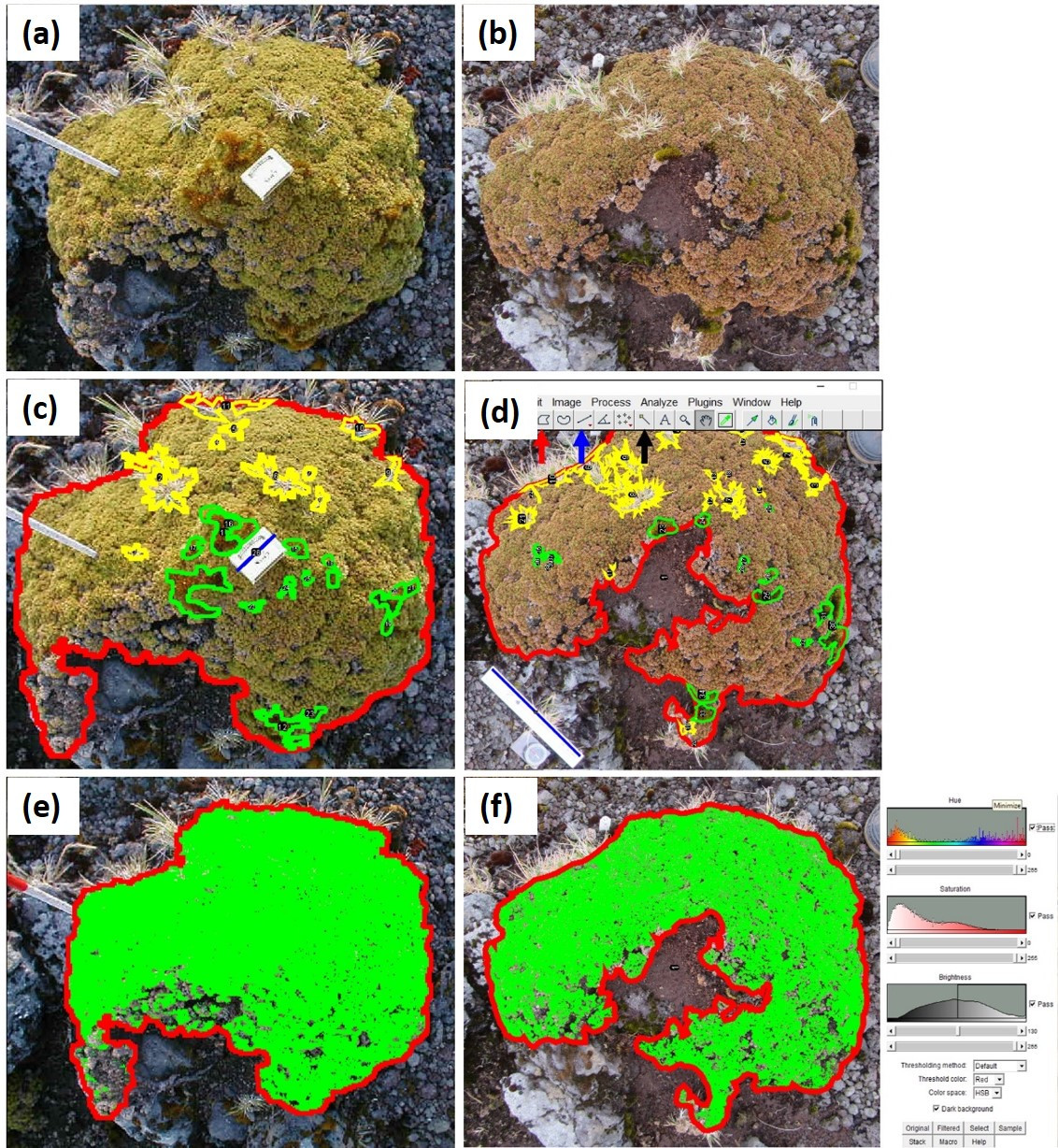

Supplementary Figure 3**.** Pictures of the same Azorella selago individual photographed in 2003 (a) and 2016 (b) respectively at a low altitude site on the western side of Marion Island. Each photograph was analysed in Image J. (c) & (d): The scale of the image was set using the length of a matchbox and ruler (blue lines and blue arrow); cushion plant circumference and area (red lines) were measured using the area selection tool (red arrow), and Agrostis magellanica area (yellow outlines) and the area of other vascular and non-vascular plants (green outlines) were measured by selecting A. magellanica individuals and other vascular and non-vascular plant individuals using the wand tracing tool (black arrow). (e) & (f): Dead stem cover on each cushion plant was measured by adjusting the colour threshold on each image (see right inset in panel (f)): green shading represents the live stem area on each cushion plant, and the remaining transparent regions are where cushion plant stems were dead after excluding the other cover types.

Supplementary Table 1. Modelling *Azorella selago* size ($n=196$), *A. selago* dead stem cover ($n=196$), and *Agrostis* *magellanica* cover ($n=163$) between 2003 and 2006 using generalized linear mixed-effects models. Cover of other = combined cover of other vascular plant species and mosses. For both categorical variables (altitude and aspect), the factors’ levels are presented according to the order of their magnitude: L = Low, M = Mid, H = High, E = East, W = West. "L vs H", "M vs H", and "W vs E", respectively, signify the difference between (i) low and high altitudes, (ii) mid and high altitudes, and (iii) western and eastern sides.

| **Response Variable** | **Statistic** | **Predictor variables: initial measurements** | | | | | | | | | | |
| --- | --- | --- | --- | --- | --- | --- | --- | --- | --- | --- | --- | --- |
|  |  | **Intercept** | ***Azorella* size (cm^2^)** | ***Agrostis* cover (%)** | **Dead stem cover (%)** | **Cover of other (%)** | **Altitude** | | **Aspect** | **% *Agrostis ×* altitude** | | **% *Agrostis ×* aspect** |
|  |  |  |  |  |  |  | **M > H > L** | | **W < E** |  |  |  |
|  |  |  |  |  |  |  | **L vs H** | **M vs H** | **W vs E** | **L vs H** | **M vs H** | **W vs E** |
| Final *Azorella* size (cm^2^) | Estimate | 1.368 | 0.866 | -1.466 | -0.002 | 0.005 | -0.051 | 0.013 | -0.024 | 1.467 | 1.466 | 0.007 |
|  | χ^2^*-*statistic | - | 2102.466 | 0.555 | 2.437 | 1.232 | 0.472 | | 0.005 | 1.211 | | 1.458 |
|  | d.f. | - | 1 | 1 | 1 | 1 | 2 | | 1 | 2 | | 1 |
|  | P-value | - | **< 0.001** | 0.456 | 0.119 | 0.267 | 0.790 | | 0.945 | 0.546 | | 0.227 |
|  | **Statistic** | **Intercept** | **Dead stem cover (%)** | ***Agrostis* cover (%)** | ***Azorella* size (cm^2^)** | **Cover of other (%)** | **Altitude** | | **Aspect** | **% *Agrostis ×* altitude** | | **% *Agrostis ×* aspect** |
|  |  |  |  |  |  |  | **L > M > H** | | **E < W** |  |  |  |
|  |  |  |  |  |  |  | **L vs H** | **M vs H** | **W vs E** | **L vs H** | **M vs H** | **W vs E** |
| Final dead stem cover (%) | Estimate | -1.090 | 0.026 | 6.058 | -0.167 | -0.016 | 0.404 | 0.014 | 0.219 | -6.049 | -6.044 | -0.026 |
|  | χ^2^-statistic | - | 67.475 | 2.106 | 9.597 | 1.375 | 0.449 | | 0.092 | 2.713 | | 2.543 |
|  | d.f. | - | 1 | 1 | 1 | 1 | 2 | | 1 | 2 | | 1 |
|  | P-value | - | **< 0.001** | 0.147 | **0.002** | 0.241 | 0.799 | | 0.762 | 0.258 | | 0.111 |
|  | **Statistic** | **Intercept** | ***Agrostis* cover (%)** | ***Azorella* size (cm^2^)** | **Dead stem cover (%)** | **Cover of other (%)** | **Altitude** | | **Aspect** | **% Dead stem *×* altitude** | | **% Dead stem *×* aspect** |
|  |  |  |  |  |  |  | **M < L** | | **W < E** |  |  |  |
|  |  |  |  |  |  |  | **M vs L** | | **W vs E** | **M vs L** | | **W vs E** |
| Final *Agrostis* cover (%) | Estimate | -3.202 | 0.058 | 0.152 | 0.002 | 0.040 | -0.903 | | -0.769 | -0.001 | | -0.007 |
|  | χ^2^-statistic | - | 116.842 | 3.866 | 0.009 | 1.631 | 22.664 | | 20.786 | 0.011 | | 0.458 |
|  | d.f. | - | 1 | 1 | 1 | 1 | 1 | | 1 | 1 | | 1 |
|  | P-value | - | **< 0.001** | **0.049** | 0.923 | 0.202 | **< 0.001** | | **< 0.001** | 0.917 | | 0.498 |

Supplementary Table 2. Modelling *Azorella selago* size ($n=194$), *A. selago* dead stem cover ($n=194$), and *Agrostis* *magellanica* cover ($n=160$) between 2006 and 2016 using generalized linear mixed-effects models. Cover of other = combined cover of other vascular plant species and mosses. For both categorical variables (altitude and aspect), the factors’ levels are presented according to the order of their magnitude: L = Low, M = Mid, H = High, E = East, W = West. "L vs H", "M vs H", and "W vs E", respectively, signify the difference between (i) low and high altitudes, (ii) mid and high altitudes, and (iii) western and eastern sides.

| **Response Variable** | **Statistic** | **Predictor variables: initial measurements** | | | | | | | | | | |
| --- | --- | --- | --- | --- | --- | --- | --- | --- | --- | --- | --- | --- |
|  |  | **Intercept** | ***Azorella* size (cm^2^)** | ***Agrostis* cover (%)** | **Dead stem cover (%)** | **Cover of other (%)** | **Altitude** | | **Aspect** | **% *Agrostis ×* altitude** | | **% *Agrostis ×* aspect** |
|  |  |  |  |  |  |  | **M > L > H** | | **E < W** |  |  |  |
|  |  |  |  |  |  |  | **L vs H** | **M vs H** | **W vs E** | **L vs H** | **M vs H** | **W vs E** |
| Final *Azorella* size (cm^2^) | Estimate | 0.510 | 0.939 | 0.000 | -0.005 | 0.011 | 0.033 | 0.064 | 0.069 | The interaction term not included due to convergence issues | | 0.004 |
|  | χ^2^-statistic | - | 1046.848 | 0.071 | 7.952 | 2.326 | 0.145 | | 0.397 |  |  | 0.383 |
|  | d.f. | - | 1 | 1 | 1 | 1 | 2 | |  |  |  | 1 |
|  | P-value | - | **< 0.001** | 0.789 | **0.005** | 0.127 | 0.930 | | 0.529 |  |  | 0.536 |
|  | **Statistic** | **Intercept** | **Dead stem cover (%)** | ***Agrostis* cover (%)** | ***Azorella* size (cm^2^)** | **Cover of other (%)** | **Altitude** | | **Aspect** | **% *Agrostis ×* altitude** | | **% *Agrostis ×* aspect** |
|  |  |  |  |  |  |  | **L > M > H** | | **E < W** |  |  |  |
|  |  |  |  |  |  |  | **L vs H** | **M vs H** | **W vs E** | **L vs H** | **M vs H** | **W vs E** |
| Final dead stem cover (%) | Estimate | -2.710 | 0.025 | -0.003 | 0.054 | 0.024 | 0.707 | 0.702 | 0.338 | The interaction term not included due to convergence issues | | -0.024 |
|  | χ^2^-statistic | - | 31.873 | 1.444 | 0.592 | 1.492 | 1.459 | | 0.189 |  |  | 2.473 |
|  | d.f. | - | 1 | 1 | 1 | 1 | 2 | | 1 |  |  | 1 |
|  | P-value | - | **< 0.001** | 0.230 | 0.442 | 0.222 | 0.482 | | 0.664 |  |  | 0.116 |
|  | **Statistic** | **Intercept** | ***Agrostis* cover (%)** | ***Azorella* size (cm^2^)** | **Dead stem cover (%)** | **Cover of other (%)** | **Altitude** | | **Aspect** | **% Dead stem *×* altitude** | | **% Dead stem *×* aspect** |
|  |  |  |  |  |  |  | **M < L** | | **W < E** |  |  |  |
|  |  |  |  |  |  |  | **M vs L** | | **W vs E** | **M vs L** | | **W vs E** |
| Final *Agrostis* cover (%) | Estimate | -3.365 | 0.050 | 0.153 | -0.008 | 0.028 | -0.569 | | -0.418 | -0.007 | | -0.005 |
|  | χ^2^-statistic | - | 115.785 | 3.418 | 4.869 | 1.044 | 5.077 | | 2.673 | 0.292 | | 0.219 |
|  | d.f. | - | 1 | 1 | 1 | 1 | 1 | | 1 | 1 | | 1 |
|  | P-value | - | **< 0.001** | 0.065 | **0.027** | 0.307 | **0.024** | | 0.102 | 0.589 | | 0.640 |

Supplementary Table 3. The number and percentage of *Azorella selago* individuals with increasing or decreasing: A) size, B) *Agrostis magellanica* cover, and C) dead stem cover, based on measurements in 2003 (i.e., initial data; indicated with subscript *i*) and in 2016 (i.e., final data; indicated with subscript *f*). Instances are indicated where *A. magellanica* cover and *A. selago* dead stem cover increased from zero initial cover (0_i_ < $x_{f}$) or increased from some initial cover ($x_{i}$ < $x_{f}$), decreased from some initial cover ($x_{i}$ > $x_{f}$) or lost all *A. magellanica* cover despite having some cover initially ($x_{i}$ > $0_{f}$).

|  | **Category** | **Number of *Azorella* individuals** | **% *Azorella* individuals** |
| --- | --- | --- | --- |
| A | *Azorella* increased in size ($x_{i}$ < $x_{f}$) | 409 | 91.9 % |
|  | *Azorella* decreased in size ($x_{i}$ > $x_{f}$) | 36 | 8.1 % |
| B | *Agrostis* cover gained ($0_{i}$ < $x_{f}$) | 34 | 7.6 % |
|  | *Agrostis* cover gained ($x_{i}$ < $x_{f}$) | 226 | 50.7 % |
|  | *Agrostis* cover lost ($x_{i}$ > $x_{f}$) | 23 | 5.2 % |
|  | *Agrostis* cover lost ($x_{i}$ > $0_{f}$) | 6 | 1.3 % |
|  | *Agrostis* absent in both years | 156 | 35.1 % |
| C | Dead stem cover increased ($x_{i}$ < $x_{f}$) | 360 | 80.9 % |
|  | Dead stem cover increased ($0_{i}$ < $x_{f}$) | 4 | 0.9 % |
|  | Dead stem cover decreased ($x_{i}$ > $x_{f}$) | 81 | 18.2 % |

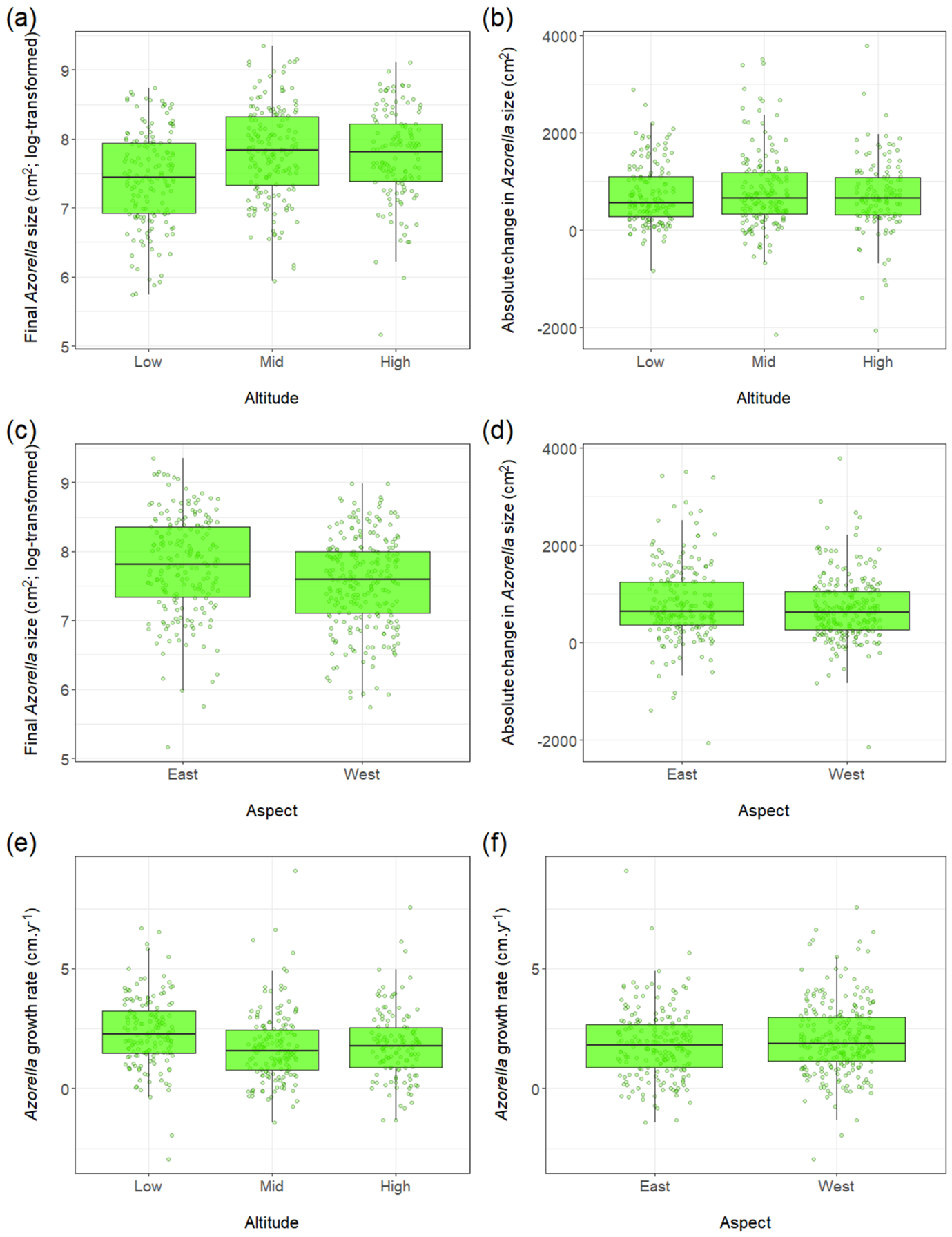

Supplementary Figure 4. Final Azorella selago size (a & c), absolute change in A. selago size (b & d) and A. selago horizontal growth rate calculated from the maximum diameter (e & f) at different altitudes and on different aspects on Marion Island. None of the differences illustrated here are significant. See Table 1 in the main text for more details.

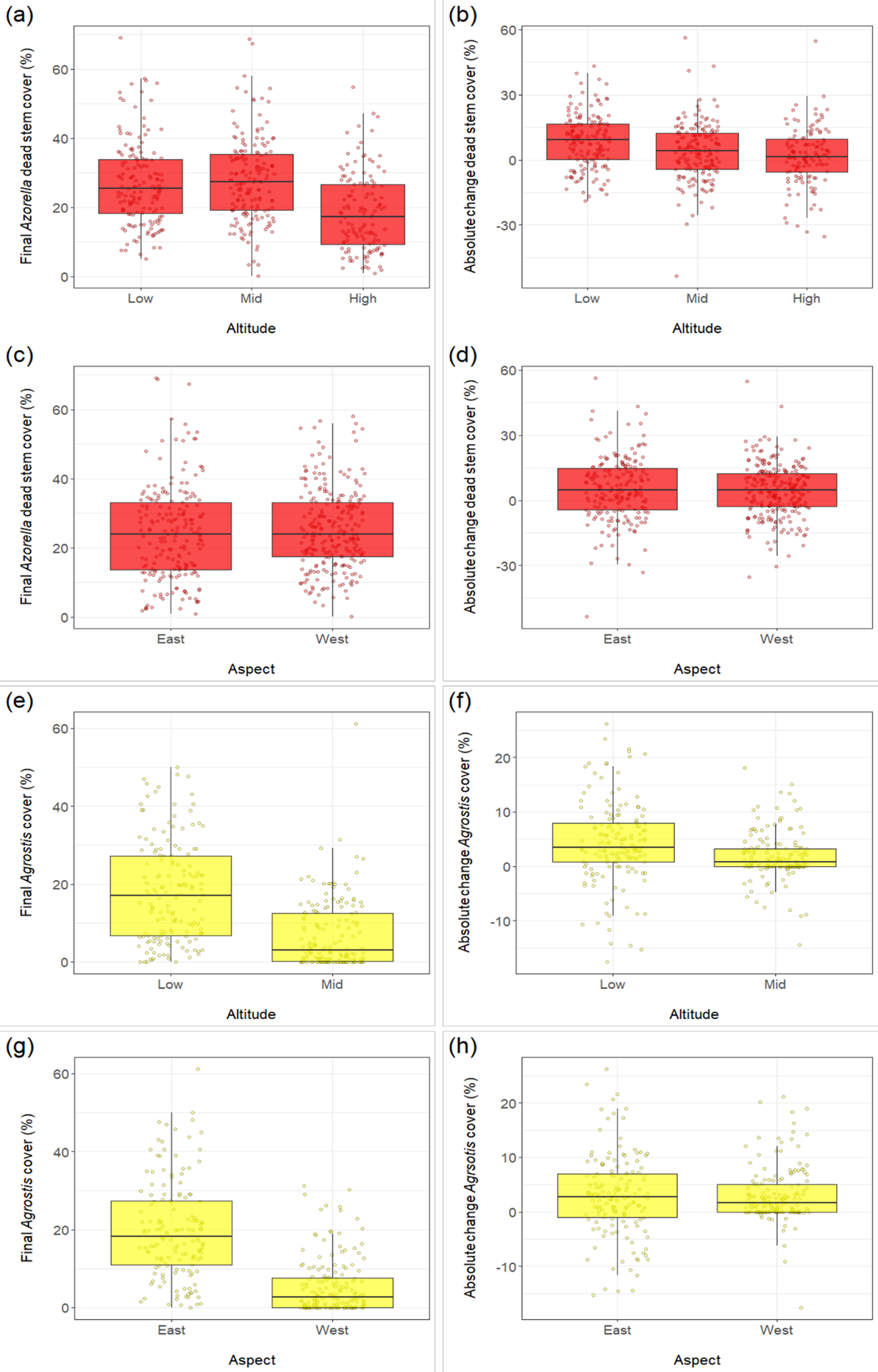

Supplementary Figure 5. Final Azorella selago dead stem cover (a & c), absolute change in A. selago dead stem cover (b & d), final Agrostis magellanica cover (e & g) and absolute change in A. magellanica cover (f & h) at different altitudes and on different aspects on Marion Island. None of the differences illustrated here are significant. See Table 1 in the main text for more details.

Supplementary Table 4. Modelling *Azorella selago* size, *A. selago* dead stem cover, and *Agrostis* *magellanica* cover separately (i.e., using a snap-shot approach) during 2003, 2006, and 2016 using generalized linear mixed-effects models. Cover of other = combined cover of other vascular plant species and mosses. For both categorical variables (altitude and aspect), the factors’ levels are presented according to the order of their magnitude: L = Low, M = Mid, H = High, E = East, W = West. "L vs H", "M vs H", and "W vs E", respectively, signify the difference between (i) low and high altitudes, (ii) mid and high altitudes, and (iii) western and eastern sides.

| **Response variable: 2003** | **Statistic** | **Predictor variables: 2003** | | | | | | | | | |
| --- | --- | --- | --- | --- | --- | --- | --- | --- | --- | --- | --- |
|  |  | **Intercept** | ***Agrostis* cover (%)** | **Dead stem cover (%)** | **Cover of other (%)** | **Altitude** | | **Aspect** | **% *Agrostis ×* altitude** | | **% *Agrostis ×* aspect** |
|  |  |  |  |  |  | **H > M > L** | | **E > W** |  |  |  |
|  |  |  |  |  |  | **L vs H** | **M vs H** | **W vs E** | **L vs H** | **M vs H** | **W vs E** |
| Final *Azorella* size (cm^2^) | Estimate | 7.606 | 0.656 | -0.004 | -0.014 | -0.687 | -0.029 | -0.201 | -0.643 | -0.636 | 0.033 |
|  | χ^2^-statistic | - | 16.418 | 2.0561 | 0.727 | 19.817 | | 0.368 | 3.719 | | 5.991 |
|  | d.f. | - | 1 | 1 | 1 | 2 | | 1 | 2 | | 1 |
|  | P-value | - | **< 0.001** | 0.152 | 0.394 | **< 0.001** | | 0.544 | 0.156 | | **0.014** |
|  | **Statistic** | **Intercept** | ***Agrostis* cover (%)** | ***Azorella* size (cm^2^)** | **Cover of other (%)** | **Altitude** | | **Aspect** | **% *Agrostis ×* altitude** | | **% *Agrostis ×* aspect** |
|  |  |  |  |  |  | **M > L > H** | | **E < W** |  |  |  |
|  |  |  |  |  |  | **L vs H** | **M vs H** | **W vs E** | **L vs H** | **M vs H** | **W vs E** |
| Final dead stem cover (%) | Estimate | -1.817 | 0.103 | -0.004 | 0.041 | 0.04 | 0.51 | 0.3 | -0.094 | -0.106 | -0.037 |
|  | χ^2^-statistic | - | 0.005 | 0.006 | 7.892 | 3.092 | | 0.548 | 1.195 | | 5.567 |
|  | d.f. | - | 1 | 1 | 1 | 2 | | 1 | 2 | | 1 |
|  | P-value | - | 0.945 | 0.939 | **0.005** | 0.213 | | 0.459 | 0.55 | | **0.018** |
|  | **Statistic** | **Intercept** | ***Azorella* size (cm^2^)** | **Dead stem cover (%)** | **Cover of other (%)** | **Altitude** | | **Aspect** | **% Dead stem *×* altitude** | | **% Dead stem *×* aspect** |
|  |  |  |  |  |  | **M < L** | | **W < E** |  |  |  |
|  |  |  |  |  |  | **M vs L** | | **W vs E** | **M vs L** | | **W vs E** |
| Final *Agrostis* cover (%) | Estimate | -3.588 | 0.328 | 0.005 | -0.003 | -1.41 | | -1.418 | 0.005 | | -0.016 |
|  | χ^2^-statistic | - | 26.819 | 0.391 | 0.008 | 53.895 | | 100.069 | 0.341 | | 3.469 |
|  | d.f. | - | 1 | 1 | 1 | 1 | | 1 | 1 | | 1 |
|  | P-value | - | **< 0.001** | 0.532 | 0.928 | **< 0.001** | | **< 0.001** | 0.559 | | 0.063 |

Supplementary Table 4 continued.

| **Response variable: 2006** | **Statistic** | **Predictor variables: 2006** | | | | | | | | | |
| --- | --- | --- | --- | --- | --- | --- | --- | --- | --- | --- | --- |
|  |  | **Intercept** | ***Agrostis* cover (%)** | **Dead stem cover (%)** | **Cover of other (%)** | **Altitude** | | **Aspect** | **% *Agrostis ×* altitude** | | **% *Agrostis ×* aspect** |
|  |  |  |  |  |  | **L < H < M** | | **E > W** |  |  |  |
|  |  |  |  |  |  | **L vs H** | **M vs H** | **W vs E** | **L vs H** | **M vs H** | **W vs E** |
| Final *Azorella* size (cm^2^) | Estimate | 7.892 | 0.009 | -0.012 | -0.001 | -0.658 | 0.025 | -0.046 | The interaction term was not included due to convergence issues | | 0.026 |
|  | χ^2^-statistic | - | 5.102 | 5.616 | 0.002 | 27.071 | | 0.266 |  |  | 4.565 |
|  | d.f. | - | 1 | 1 | 1 | 2 | | 1 |  |  | 1 |
|  | P-value | - | **0.024** | **0.018** | 0.964 | **< 0.001** | | 0.606 |  |  | **0.033** |
|  | **Statistic** | **Intercept** | ***Agrostis* cover (%)** | ***Azorella* size (cm^2^)** | **Cover of other (%)** | **Altitude** | | **Aspect** | **% *Agrostis ×* altitude** | | **% *Agrostis ×* aspect** |
|  |  |  |  |  |  | **H < M < L** | | **W > E** |  |  |  |
|  |  |  |  |  |  | **L vs H** | **M vs H** | **W vs E** | **L vs H** | **M vs H** | **W vs E** |
| Final dead stem cover (%) | Estimate | -0.807 | 0.012 | -0.152 | 0.007 | 0.481 | 0.255 | 0.244 | The interaction term was not included due to convergence issues | | -0.028 |
|  | χ^2^-statistic | - | 1.736 | 5.348 | 0.196 | 0.469 | | 0.036 |  |  | 4.473 |
|  | d.f. | - | 1 | 1 | 1 | 2 | | 1 |  |  | 1 |
|  | P-value | - | 0.188 | **0.021** | 0.658 | 0.791 | | 0.850 |  |  | **0.034** |
|  | **Statistic** | **Intercept** | ***Azorella* size (cm^2^)** | **Dead stem cover (%)** | **Cover of other (%)** | **Altitude** | | **Aspect** | **% Dead stem *×* altitude** | | **% Dead stem *×* aspect** |
|  |  |  |  |  |  | **M < L** | | **W < E** |  |  |  |
|  |  |  |  |  |  | **M vs L** | | **W vs E** | **M vs L** | | **W vs E** |
| Final *Agrostis* cover (%) | Estimate | -3.318 | 0.312 | 0.009 | 0.021 | -1.826 | | -0.945 | 0.020 | | -0.028 |
|  | χ^2^-statistic | - | 9.859 | 1.465 | 0.312 | 101.181 | | 109.162 | 1.936 | | 5.317 |
|  | d.f. | - | 1 | 1 | 1 | 1 | | 1 | 1 | | 1 |
|  | P-value | - | **0.002** | 0.226 | 0.576 | **< 0.001** | | **< 0.001** | 0.164 | | **0.021** |

Supplementary Table 4 continued.

| **Response variable: 2016** | **Statistic** | **Predictor variables: 2016** | | | | | | | | | |
| --- | --- | --- | --- | --- | --- | --- | --- | --- | --- | --- | --- |
|  |  | **Intercept** | ***Agrostis* cover (%)** | **Dead stem cover (%)** | **Cover of other (%)** | **Altitude** | | **Aspect** | **% *Agrostis ×* altitude** | | **% *Agrostis ×* aspect** |
|  |  |  |  |  |  | **M > H > L** | | **W < E** |  |  |  |
|  |  |  |  |  |  | **L vs H** | **M vs H** | **W vs E** | **L vs H** | **M vs H** | **W vs E** |
| Final *Azorella* size (cm^2^) | Estimate | 8.067 | 0.292 | -0.008 | -0.034 | -0.554 | 0.001 | -0.172 | -0.279 | -0.277 | 0.022 |
|  | χ^2^-statistic | - | 18.153 | 10.392 | 10.635 | 17.506 | | 0.099 | 2.017 | | 6.128 |
|  | d.f. | - | 1 | 1 | 1 | 2 | | 1 | 2 | | 1 |
|  | P-value | - | **< 0.001** | **0.001** | **0.001** | **< 0.001** | | 0.753 | 0.365 | | **0.013** |
|  | **Statistic** | **Intercept** | ***Agrostis* cover (%)** | ***Azorella* size (cm^2^)** | **Cover of other (%)** | **Altitude** | | **Aspect** | **% *Agrostis ×* altitude** | | **% *Agrostis ×* aspect** |
|  |  |  |  |  |  | **M > L > H** | | **E < W** |  |  |  |
|  |  |  |  |  |  | **L vs H** | **M vs H** | **W vs E** | **L vs H** | **M vs H** | **W vs E** |
| Final dead stem cover (%) | Estimate | -0.881 | 0.328 | -0.1060 | 0.005 | 0.586 | 0.607 | 0.222 | -0.326 | -0.325 | -0.015 |
|  | χ^2^-statistic | - | 0.042 | 5.299 | 0.282 | 12.613 | | 0.787 | 3.115 | | 1.966 |
|  | d.f. | - | 1 | 1 | 1 | 2 | | 1 | 2 | | 1 |
|  | P-value | - | 0.838 | **0.021** | 0.596 | **0.002** | | 0.375 | 0.211 | | **0.161** |
|  | **Statistic** | **Intercept** | ***Azorella* size (cm^2^)** | **Dead stem cover (%)** | **Cover of other (%)** | **Altitude** | | **Aspect** | **% Dead stem *×* altitude** | | **% Dead stem *×* aspect** |
|  |  |  |  |  |  | **M < L** | | **W < E** |  |  |  |
|  |  |  |  |  |  | **M vs L** | | **W vs E** | **M vs L** | | **W vs E** |
| Final *Agrostis* cover (%) | Estimate | -3.192 | 0.317 | -0.007 | 0.048 | -1.396 | | -1.370 | 0.005 | | -0.005 |
|  | χ^2^-statistic | - | 24.772 | 3.670 | 16.516 | 63.071 | | 88.460 | 0.543 | | 0.553 |
|  | d.f. | - | 1 | 1 | 1 | 1 | | 1 | 1 | | 1 |
|  | P-value | - | **< 0.001** | 0.057 | **< 0.001** | **< 0.001** | | **< 0.001** | 0.461 | | 0.457 |

Supplementary Table 5. Modelling the number of fruits ($n=214$) and flower buds ($n=214$) on *Azorella selago* during 2003 using generalized linear mixed-effects models. Cover of other = combined cover of other vascular plant species and mosses. For both categorical variables (altitude and aspect), the factors’ levels are presented according to the order of their magnitude: L = Low, M = Mid, H = High, E = East, W = West. "L vs H", "M vs H", and "W vs E", respectively, signify the difference between (i) low and high altitudes, (ii) mid and high altitudes, and (iii) western and eastern sides. *Azorella* size (cm^2^) has been included as an offset variable.

| **Response variable: 2003** | **Statistic** | **Predictor variables: 2003** | | | | | | | | |
| --- | --- | --- | --- | --- | --- | --- | --- | --- | --- | --- |
|  |  | **Intercept** | ***Agrostis* cover (%)** | **Cover of other (%)** | **Altitude** | | **Aspect** | **% *Agrostis ×* altitude** | | **% *Agrostis ×* aspect** |
|  |  |  |  |  | **L < M < H** | | **W < E** |  |  |  |
|  |  |  |  |  | **L vs H** | **M vs H** | **W vs E** | **L vs H** | **M vs H** | **W vs E** |
| Number of fruits | Estimate | -0.617 | 1.013 | -0.102 | -0.902 | -0.728 | -0.355 | -1.032 | -1.037 | -0.057 |
|  | χ^2^-statistic | - | 5.378 | 1.244 | 3.502 | | 3.782 | 0.373 | | 1.506 |
|  | d.f. | - | 1 | 1 | 2 | | 1 | 2 | | 1 |
|  | P-value | - | **0.020** | 0.265 | 0.174 | | 0.052 | 0.830 | | 0.220 |
|  | **Statistic** | **Intercept** | ***Agrostis* cover (%)** | **Cover of other (%)** | **Altitude** | | **Aspect** | **% *Agrostis ×* altitude** | | **% *Agrostis ×* aspect** |
|  |  |  |  |  | **H > M > L** | | **E < W** |  |  |  |
|  |  |  |  |  | **L vs H** | **M vs H** | **W vs E** | **L vs H** | **M vs H** | **W vs E** |
| Number of flower buds | Estimate | -1.505 | -2.873 | -0.056 | -1.666 | -0.761 | 1.126 | 2.849 | 2.832 | -0.030 |
|  | χ^2^-statistic | - | 6.089 | 0.763 | 10.351 | | 7.490 | 3.0276 | | 1.152 |
|  | d.f. | - | 1 | 1 | 2 | | 1 | 2 | | 1 |
|  | P-value | - | **0.014** | 0.382 | **0.006** | | **0.006** | 0.220 | | 0.283 |

Supplementary Text 1

Summary of the methodology utilized by Nyakatya (2006)

The research by Nyakatya (2006) has been published as MSc thesis (available from: <https://scholar.sun.ac.za/handle/10019.1/21696>) but here I summarize the key study design points relevant to the methods used for our study. The broad aim of Nyakatya’s (2006) study was to quantify spatial variability in the phenology, morphology, reproductive effort, and epiphyte load of *Azorella selago* cushion plants across sub-Antarctic Marion Island, and to determine the direction and range of this variability.

Twelve long-term monitoring plots were established and surveyed at three altitudes (c. 200, 400 and 600 m a.s.l) on the island’s eastern and western aspects between April 2002 and April 2003. Plots were established using complete sampling, i.e., a central starting point was selected, and the area encompassing a minimum of 50 *A. selago* plants (excluding individuals < 15 cm diameter) from that starting point was considered as a plot. The exact locations of each plot were randomly selected within certain constraints: 1) plots had to be in *Azorella*-dominated fellfield; 2) plots had to be located within defined altitudinal bands (i.e., 150 - 250 m a.s.l., 350 - 450 m a.s.l. and > 550 m a.s.l.); and 3) plots needed to form an altitudinal transect (two altitudinal transects were established on both the eastern and western sectors of the island). The plots were clearly marked with corner marker poles and tags for long-term monitoring purposes. Within each plot, 50 *A. selago* cushion plants (greater than 15 cm in diameter) were selected and used for taking several of measurements. Non-destructive measurements were also taken from cushion plants that were less than 15 cm in diameter to avoid damaging young plants. The exact and relative position of each cushion plant within a site, its nearest neighbours, and the corners of each sampling site were determined using a Nikon Total Station DTM350 Theodolite, with an accuracy of 10 mm. Since there are no fixed reference points on the island, a Garmin 12MAP GPS (global positioning systems) was used to obtain the approximate geographic co-ordinates of each site.

Within these sites, a variety of quantitative measurements of *A. selago* were recorded*.* Each of the 50 *A. selago* individual within each site was photographed in the summer of 2002/2003 from directly above at a height of 1.5 m, with a scale bar included within each photograph.

Supplementary Text 2

The statistical models specified for our analyses:

1. *log(final Azorella size (cm^2^)) ~ log(initial Azorella size (cm^2^)) + initial Agrostis cover (%) + initial Azorella dead stem cover (%) + altitude + aspect + other vascular plants and mosses’ initial combined cover (%) + (initial Agrostis cover (%) × altitude) + (initial Agrostis cover (%) × aspect) + (1│plot)*	***(Eqn. 1)***
2. *Final Azorella dead stem cover (%) ~ initial Azorella dead stem cover (%) + log(initial Azorella size (cm^2^)) + initial Agrostis cover (%) + altitude + aspect + other vascular plants and mosses’ initial combined cover (%) + (initial Agrostis cover (%) × altitude) + (initial Agrostis cover (%) × aspect) + (1│plot)*	***(Eqn. 2)***
3. *Final Agrostis cover (%) ~ initial Agrostis cover (%) + log(initial Azorella size (cm^2^)) + initial Azorella dead stem cover (%) + altitude + aspect + other vascular plants and mosses’ combined cover (%) + (initial dead stem cover (%) × altitude) + (initial dead stem cover (%) × aspect) + (1│plot)*	***(Eqn. 3)***
